# Sequencing of distinct wing behaviors during *Drosophila* courtship

**DOI:** 10.1101/2025.08.21.671456

**Authors:** Xinping Li, Kyle Thieringer, Yiqin Gao, Mala Murthy

## Abstract

Some behaviors, like biting followed by chewing and then swallowing, unfold in stereotyped sequences, while others, such as limb movements during defensive maneuvers, can be flexibly combined as needed. During courtship, male *Drosophilid* flies produce a series of actions, including orientation, tapping, singing, licking, and copulation, that follow an ordered but temporally variable sequence [1,2]. At shorter timescales, however, individual actions remain highly dynamic. For example, courtship songs are composed of variable sequences of distinct syllables, with their patterning and amplitude actively shaped by female cues [3–5]. Leveraging recent advances in behavioral quantification [6], we discover a new courtship wing behavior that we term “waggling”, which is present across multiple *Drosophila* species and characterized by rhythmic, anti-phase wing movements. We identify an intermediate level of stereotyped behavioral structure: a directional three-part motif where males and females first decelerate to near-complete stillness, followed by male-initiated waggling, which then transitions into courtship song. Wing kinematics during waggle bouts are predictive of wing choice in subsequent songs, suggesting waggling may serve as a preparatory behavior. We then focus on P1/pC1 neurons, known to promote courtship [5,7–11]. Optogenetic activation of specific P1/pC1 neuron subsets in solitary males, without any female cues, is sufficient to recapitulate the entire stillness-to-waggling-to-singing progression. These findings reveal a new layer of stereotyped structure within a flexible courtship display and demonstrate that P1/pC1 neurons can orchestrate multi-action behavioral programs through internal dynamics.

**Highlights:**

- *Drosophila* males produce “waggling,” an oscillatory, anti-phase movement of the two wings that is distinct from the unilateral wing vibration that generates courtship song.
- Waggling is part of a structured behavioral sequence: from stillness, to waggling, and then to singing during male-female courtship interactions.
- This full behavioral sequence is internally driven and can be triggered in solitary males by optogenetic activation of specific subsets of P1/pC1 neurons.
- P1/pC1 neurons show functional diversity in locomotor control.

## Results and Discussion

### *Drosophila* males produce two distinct wing behaviors during courtship -singing and waggling

We used quantitative pose estimation (via SLEAP [6]) of a large dataset of *Drosophila melanogaster* male-female interactions (32 pairs, 13.4 hours of videos), along with audio recordings, to identify a previously undescribed non-acoustic wing behavior during fly courtship, which we term wing “waggling”. During waggling, *D. melanogaster* males extend both wings (at reduced angles relative to song [12]), in contrast to the characteristic unilateral wing extension of singing (Fig. 1A-B). The defining feature of waggling is the rapid, anti-phase oscillation of the wings at a median frequency of approximately 10.6 Hz (Fig. 1C), resembling the motion of windshield wipers (Videos S1-2). This frequency is intermediate between the recently described low-frequency thoracic vibrations [13,14] and the higher-frequency wing vibrations that generate courtship song [12,15,16]. While previous studies have described a “wing scissoring” behavior [15,17–20], scissoring has never been quantitatively defined. In fact, this term appears to have been applied to various wing displays, including wing waving, bilateral wing spreading [21], and potential behaviors that may overlap with what we term waggling. For this reason, we use a distinct terminology and define waggling as an oscillatory behavior of the two wings.

**Figure 1.**
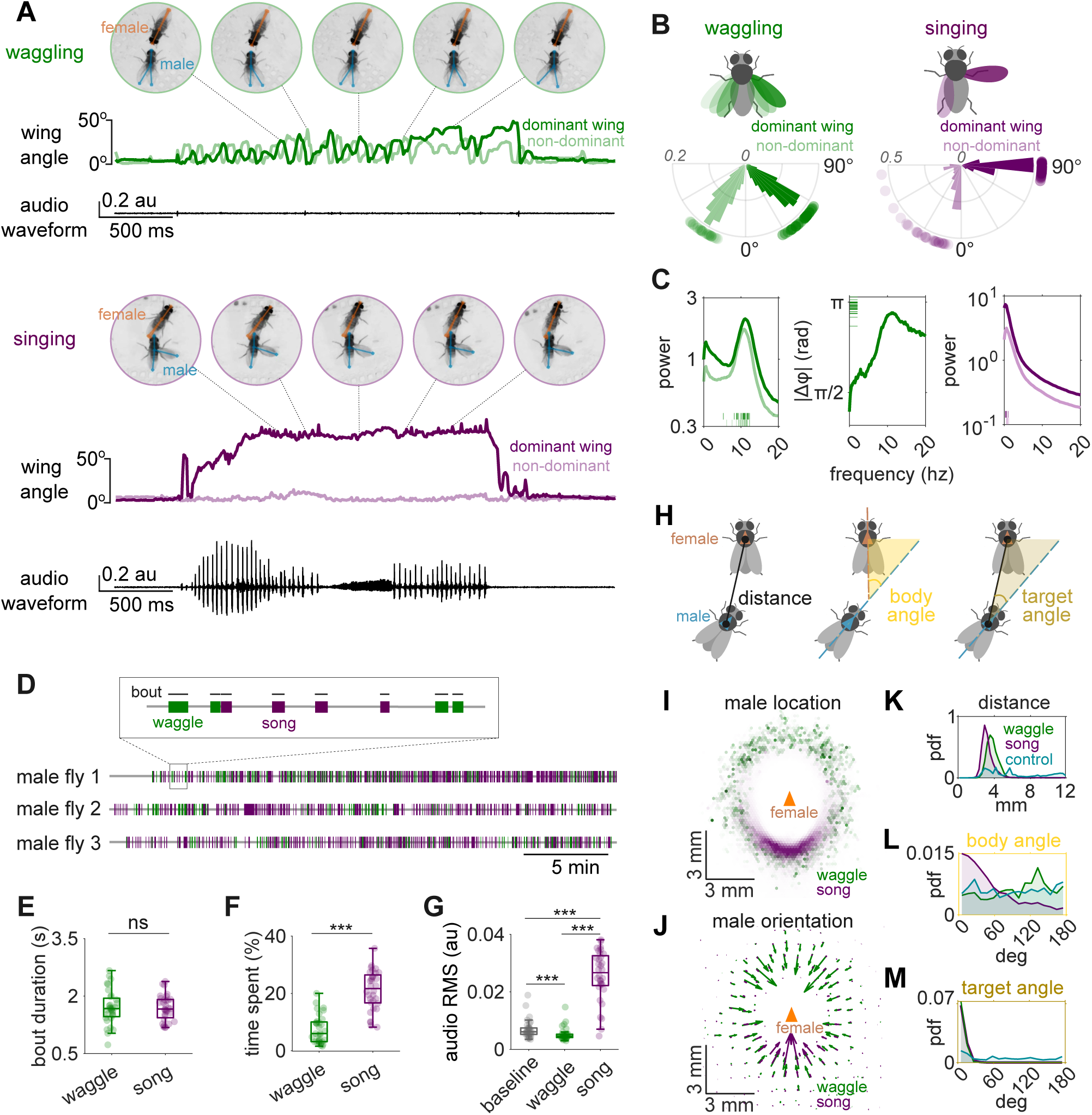
*Drosophila melanogaster* males produce two distinct wing behaviors during courtship. A. An example waggling bout (top) and courtship song bout (bottom). Snapshots (with SLEAP^6^ skeletons overlaid) indicate discrete moments in each behavior during male/female interactions. Male wing extension angles are different during waggling versus singing (the dominant wing is defined as the wing with greater mean amplitude throughout the bout); waggling involves an oscillatory behavior between the two wings. Waggling does not produce a detectable sound on the microphones (see Methods). B. Wing extension angles. Polar histograms plot the distribution of maximum wing extension angles for waggling (green, 1 857 bouts) and singing (purple, 5087 bouts), the radial axis shows probability density, and dots outside each plot mark the per-fly medians of these peak angles. C. Frequency-domain analysis. Left two panels (green): Mean power spectra (± SEM) of dominant and non-dominant wings during waggling (left) and mean absolute dominant/non-dominant wing phase difference (right); insets show the distribution of each individual fly’s median values. Right-most panel (purple): Mean power spectra (± SEM) of dominant and non-dominant wings during singing. D. Gantt-style timelines plot all waggling (green) and singing (purple) bouts detected in three representative recordings; each horizontal line is one fly pair, and colored bars mark the start-to-end time of individual waggle or song bouts (see Methods). Waggling and singing both occur throughout courtship. E. Waggle and song bout durations are comparable (paired t-test, p=0.901; ns, not significant). F. Males spend a larger fraction of the recording singing versus waggling (Wilcoxon signed-rank test, *** p < 0.001). G. RMS (Root Mean Square) microphone amplitude is highest during song bouts, intermediate at baseline, and lowest during waggling bouts (Friedman test with Wilcoxon post-hoc, *** p < 0.001). H. Spatial geometry of the courting male relative to the female (see Methods). I. Female-centered map of male location during waggling (green) or singing (purple). The orange triangle marks the female thorax and her orientation. Males stay close in both behaviours but, whereas singing occurs mostly directly behind the female, waggling is produced all around the female. J. Female-centered quiver plot of male orientation during waggling (green) or singing (purple). Males orient towards the female during both behaviors. K-M. Distributions of male–female spatial parameters (see H). Probability-density functions show (K) distance, (L) body angle, and (M) target angle during waggling (green), singing (purple) and control (cyan; frames that precede the first waggle or song frame in each recording). N=32 males across panels B-C and E-M. For E-G, box plots show median ± IQR, whiskers extend to 1.5×IQR. Dots mark the individual per-fly medians.

Similar to singing, waggling is never observed in solitary males and occurs in bouts throughout male-female interactions. While waggling occurs less frequently than singing under our recording conditions, it represents a significant component of male courtship behavior (Fig. 1D-F). However, waggling does not generate an audible signal (Fig. 1G). These properties are conserved across species. Using similar methods for behavioral quantification, we observed waggling in both *D. simulans* and *D. sechellia* (Videos S3-4, Fig. S1); both species are separated from the *D. melanogaster* lineage only about 3 million years ago [22]. While *D. sechellia* exhibited greater wing extension angles and lower oscillation frequencies compared to *D. melanogaster* and *D. simulans*, the presence of anti-phase oscillatory wing movements across species highlights that waggling is likely an important courtship behavior.

While we do not know if waggling serves a communicative purpose, it only occurs when males are close to females (within 3-4 mm) and are oriented toward them (target angle ∼0°), suggesting it may be driven by cues from the female. One key difference with singing is that while song is produced mostly behind the female (often while chasing her [3]), waggling is produced all around the female (Fig. 1H-M). Below we investigate the coordination between singing and waggling.

### Waggling is part of a behavioral sequence during courtship

Song production is linked to locomotor dynamics - male and female speed can predict both the timing and patterning of song [3]. Is the same true for waggling? We found that waggling and singing showed distinct, and often opposite, associated locomotor dynamics (Fig. 2A-B). The onset of a waggling bout is preceded by marked deceleration of both males and females, who both remain stationary prior to and throughout the behavior (Fig. 2C-E, Fig. S2). This contrasts with singing, which is typically marked by acceleration prior to a song bout, as well as continued motion of both male and female throughout the song bout. These findings further establish waggling as a distinct behavioral state with specific locomotor requirements. It is worth noting that the stillness phase (when both males and females remain stationary) can potentially overlap with abdominal vibrations, which are used as a communication signal [13,14], but not measured here.

**Figure 2.**
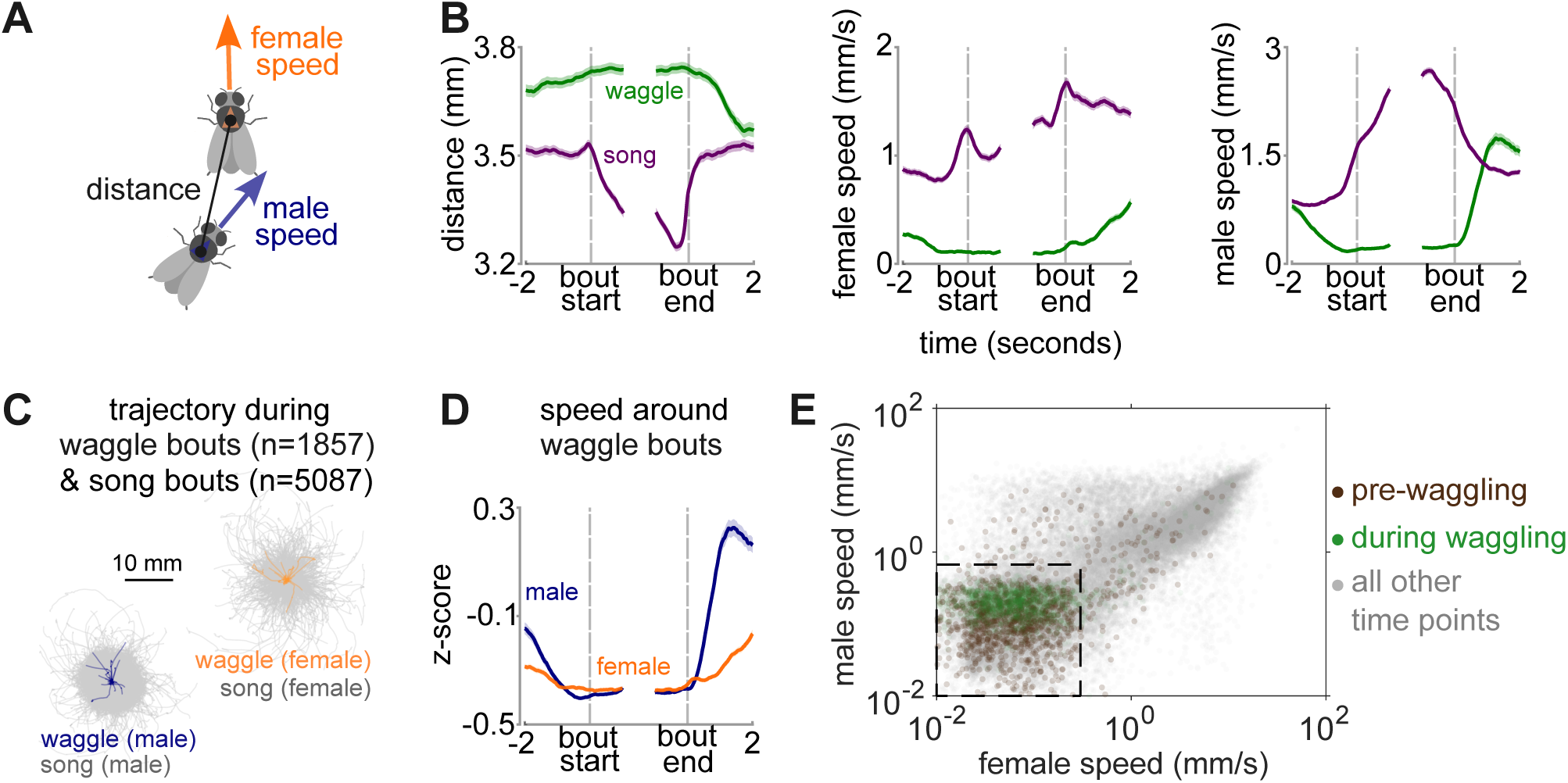
Locomotor dynamics differ around waggle versus song bouts. A. Parameters analyzed: the distance between the male and female thorax, the speed of the male or female, along with their heading directions. B. Bout-triggered averages (mean ± SEM) of male–female distance (left), female speed (middle), and male speed (right) aligned to bout start and end. Green: waggle bouts; purple: song bouts. Waggle occurs at larger male-female distances and slower speeds relative to song. C. Male and female trajectories during individual waggle (male: navy; female: orange) and song bouts (gray). All trajectories are centered on the bout start position. D. Z-scored speeds around waggle bouts. Bout-triggered averages after z-scoring speeds within individual animal recordings (same data as B but z-scored). E. Scatter plot of male versus female speeds during pre-waggling (brown, 1 s before bouts), waggling (green, entire bout averages), and all other times (gray, 1 s bins). Boxed region marks the “stillness” zone (0 -- 0.67 mm/s male, 0 -- 0.3 mm/s female; defined in Figure S2). N=32 male-female pairs in B-E.

In addition, we observed that waggling often precedes song but not the other way around (Fig. S3, also see Fig. 4A-B). We therefore examined the relationship between waggle bouts and the song bouts that follow. We found that waggling bouts often transition directly into song bouts, a sequence we term a “linked” waggle (Fig. 3A). These linked transitions have distinct dynamic signatures compared to “unlinked” waggles that are not immediately followed by song. Specifically, these linked bouts are associated with closer male-female proximity, lower female speed both before and after waggle offset, and higher male speed immediately following the waggle (Fig. 3B-C). Furthermore, wing choice for the subsequent song bout is strongly correlated with the dominant wing used during the preceding waggle, especially in linked transitions (Fig. 3D). The shorter the time gap between waggling and singing, the stronger the correlation. Remarkably, this coupling effect persists for over 14 seconds before eventually decaying back to chance levels (Fig. 3D). These findings suggest that waggling may serve as a preparatory behavior that facilitates wing extension and selection during subsequent singing.

**Figure 3.**
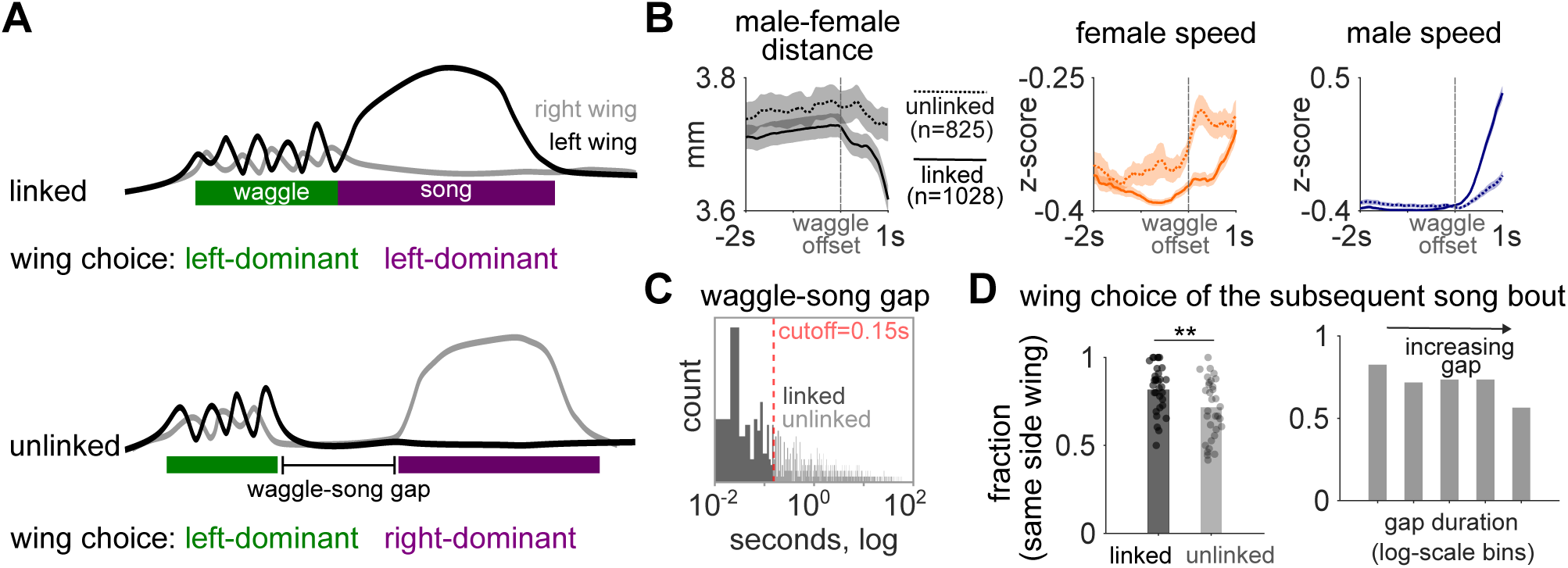
Waggling is linked to singing. A. A waggle bout can either be directly followed by song (top: “linked” waggle) or not (bottom: “unlinked” waggle). In either case, males can use the same wing as dominant during waggling and subsequent singing (shown at top) or they can switch the dominant wing between the two behaviors (shown at bottom). B. Locomotor dynamics around waggle bout ends for linked (solid lines) and unlinked (dashed lines) transitions to song. Bout-triggered averages (mean ± SEM) of male–female distance (left) and speeds (female: middle; male: right) aligned to waggle bout ends. Linked waggles involve a steep drop in male-female distance as males transition from waggling to singing, as well as a sharp increase in male and female speeds. C. Distribution of gap durations between the end of waggle bouts and the start of the subsequent song bout. The red dashed line marks the cutoff (0.15 s) separating linked and unlinked transitions (see Methods). D. Wing choice in song bouts following waggling. Left panel: Fraction of song bouts where the dominant wing matches the dominant wing of the preceding waggle bout for unlinked and linked transitions. Dots mark individual animals (paired t-test, **p < 0.01). Right panel: Same data binned by waggle–song gap duration (log-scale bins: 0.007–0.046 s, n=947; 0.046–0.312 s, n=145; 0.312–2.138 s, n=290; 2.138–14.630 s, n=393; 14.630–100.133 s, n=78). The more linked the waggle and song bouts are, the more likely the dominant wing will be the same for the two behaviors. N=32 males in B-D.

**Figure 4.**
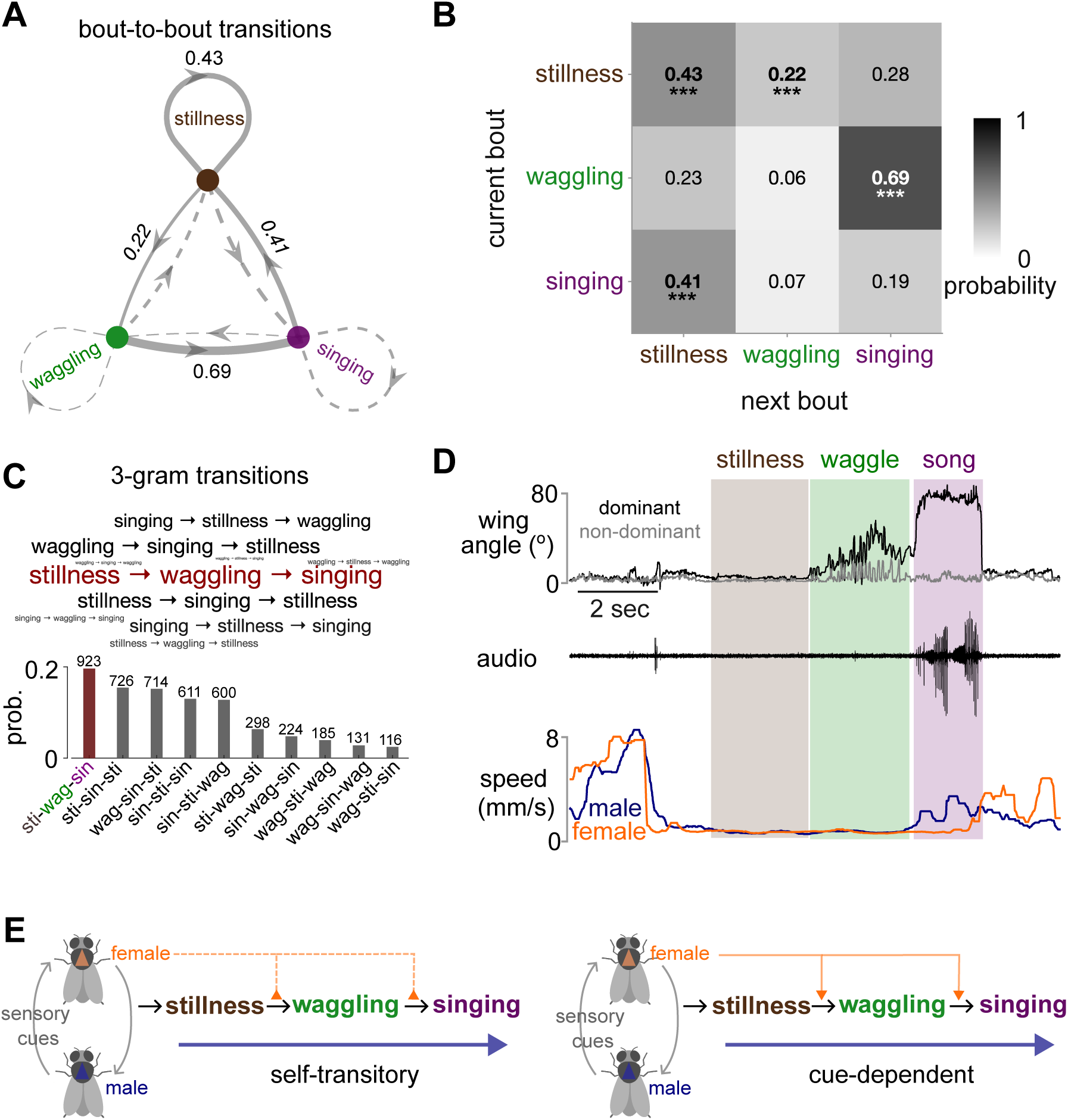
Sequencing of stillness, waggling, and singing. A-B. Bout-to-bout behavioral state transitions. Network diagram (A) and transition probability matrix (B) showing probabilities of transitioning between three different kinds of ‘bouts’: stillness, waggling, and singing. Only three states are shown for clarity (see Figure S4 for full four-state matrix that includes “other” for all other behaviors produced during courtship); probabilities do not sum to 1 as transitions to the “other” state are excluded. Solid arrows and asterisks indicate statistically significant transitions (one-tailed permutation test, ***p < 0.001). See Methods for stillness, waggling and singing bout definitions. C. Three-gram transition probabilities. Bars show probability of each three-bout sequence, with counts above. Word cloud shows the most frequent three-gram sequences, with font size scaled to frequency. D. Representative sequence from stillness to waggling and to singing. Wing angles (dominant in black and non-dominant in gray), audio recordings, and male (blue) and female (orange) speeds during a single recording (see Video S5). E. Conceptual models of behavioral sequencing during courtship. Left: Self-transitory model: male’s internal dynamics drive behavioral transitions, with female sensory cues modulating the probability of staying in or exiting each state. Right: Cue-dependent model: female sensory cues directly drive transitions from one behavior to the next. N=32 male-female pairs in A-C.

We next analyzed behavioral transitions between stillness (periods when both males and females remain stationary), waggling, and singing, which revealed a clear, directional structure (Fig. 4 A-B, Fig. S4). The transition probability from waggling to singing is exceptionally high, and the most frequent three-state sequence was stillness → waggling → singing (Fig. 4C-D, Video S5). This progression suggests a stereotyped behavioral sequence within courtship interactions.

Two mechanistic models, both supported by existing literature [3,23], could explain this behavioral sequence (Fig. 4E): a “self-transitory” model where male internal dynamics drive the sequence progression with female cues possibly gating the transitions, or a “cue-dependent” model where female sensory inputs directly drive each behavior in the sequence. Distinguishing these requires examining behavior in the absence of female cues.

### P1 neuron subset orchestrates the behavioral sequence by modulating locomotion

We next used optogenetics to uncover the cell types that promote the waggling and the courtship sequence described above (Fig. 5A-B). P1/pC1 neurons are a sexually dimorphic group of roughly 60 neurons per hemisphere that are capable of driving a persistent arousal state [5,7,24,25] and that promote a variety of social behaviors, including both courtship and aggressive actions [8,10,26–28]. For simplicity, we will refer to P1/pC1 as P1 throughout the text. Within this population, activation of a male-specific subset of 6-8 P1 neurons per hemisphere (called P1a) can drive acute locomotor arrest [12,13,24]. In addition, P1a activation, in solitary male flies, does not drive courtship singing directly, but rather a persistent state of elevated song production [5]. Because of this, we hypothesized that P1 neurons might orchestrate the behavioral program of stillness leading to waggling and then singing (Fig. 4).

**Figure 5.**
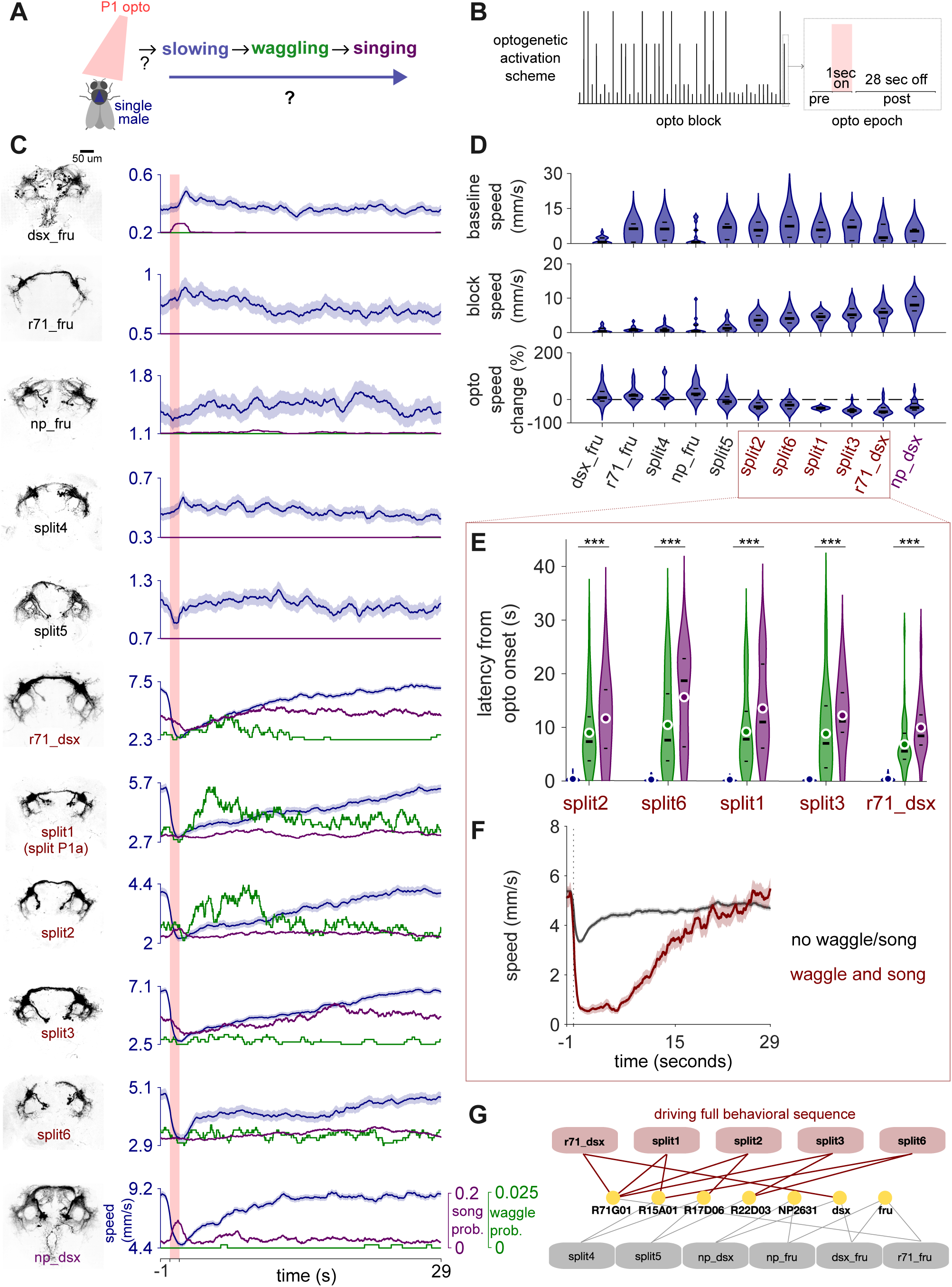
P1 neuronal subsets drive behavioral sequences that include waggling. A. Experimental design. Optogenetic activation of P1 neuronal subsets to investigate the production of slowing, waggling, and singing states in the absence of female cues. B. Optogenetic stimulation protocol (see Methods). A one second light pulse (of varying amplitude) delivered every 30 seconds across the stimulation block. C. Behavioral responses across 11 different genotypes targeting different P1 neuronal subsets. Left: Anatomical expression patterns. Right: Mean ± SEM walking speed (blue) aligned to LED light onset (red bar), with waggling (green) and singing (purple) probabilities. Genotypes with red labels exhibit robust activation of the full behavioral sequence (slowing → waggling → singing). D. Summary of optogenetically-induced speed responses across genotypes. Violin plots show distribution across animals for baseline speed (top; prior to the start of optogenetic stimulation), block speed (middle; speed during the entire optogenetic block), and percentage change (bottom; calculated as (during-pre)/pre, comparing speed during optogenetic stimulation (when red LED is on) to pre-stimulation baseline). E. Latency from optogenetic onset to behavioral responses in genotypes that drive the full behavioral sequence. Violin plots show distribution across trials for peak deceleration (blue), waggling onset (green), and singing onset (purple). Friedman test within each genotype (***p < 0.001). F. Optogenetic epoch-triggered averages (mean ± SEM) aligned to optogenetic onset in genotypes driving the full behavioral sequence, split by whether stimulation evoked (red) or did not evoke (black) the behavioral sequence. G. Network of genetic driver combinations for P1 neurons and functional outcomes. Connections indicate genetic driver combinations; those producing full behavioral sequences are indicated in red, others in gray. See Table S1 for full information on genetic driver lines. N = 17 (dsx_fru), 15 (np_dsx), 14 (np_fru), 19 (r71_dsx), 15 (r71_fru), 19 (split1/split P1a), 18 (split2), 19 (split3), 19 (split4), 19 (split5), 18 (split6) males. Violin plots show median and quartiles (black lines, in D and E) and means (white circles, in E).

P1 neurons contain both Doublesex and Fruitless expressing subsets [28–30], comprising multiple genetically and functionally heterogeneous subtypes [8,31–34]. While most studies of P1 have so far focused on P1a, we instead selected and tested multiple genetic lines known to label broader groupings of P1 neurons [24,33,34] (Fig. 5C, Fig. S5, Table S1). These lines each target a different subset of P1 neurons. By activating these P1 sub-populations in solitary males (without the presence of female sensory cues), we could test whether internal neural dynamics alone can drive the behavioral progression from stillness to waggling to singing (Fig. 5A). We chose an optogenetic paradigm in which we interleaved brief (1 second) stimuli with long intervals (Fig. 5B), in order to uncover behavioral sequences that followed stimulation - this is in contrast with prior work that used higher duty cycles for P1 neuron activation [5,12].

Screening of 11 genetically defined P1 subsets revealed two distinct response classes based on acute locomotor effects (Fig. 5C-D). Activation of some P1 subsets (split2, split 6, split1 (also known as P1a), split3, r71_dsx, np_dsx) showed acute deceleration during optogenetic stimulation but elevated overall locomotor speed, while the other group (dsx_fru, r71_fru, split4, np_fru, split5) showed either acceleration or no change during acute stimulation but exhibited suppressed overall locomotor speed. In addition, the acute locomotor effects, regardless of direction, were dose-dependent and influenced by the initial locomotor state of the fly at stimulation onset (Fig. S6). Only drivers inducing acute slowing (r71_dsx, split1/P1a, split2, split3, split6) reliably triggered the complete waggling to song behavioral sequence (Fig. 5C-E, Fig. S7). The kinematics of this optogenetically-evoked waggling, including its anti-phase wing movements, closely matched those of the natural behavior (Fig. S8). The latency from optogenetic onset to peak slowing, then waggling, then singing followed a consistent temporal order that mirrored the natural progression (Fig. 5E). In addition, locomotor state modulated sequence probability—flies that slowed down sufficiently were more likely to complete the sequence, suggesting locomotion itself acts as an internal context that shapes behavioral state transitions (Fig. 5F). For instance, np_dsx P1 activation drove very little waggling and singing following optogenetic activation - males of this genotype had a higher overall speed, and so the slowing induced by optogenetic activation did not cause enough of a reduction in speed to reliably lead to the subsequent behaviors (Fig. 5C).

Taken together, our results strongly support the self-transitory model (Fig. 4E): external sensory input is not needed to transition from stillness to waggling to singing; instead P1 neurons (and specifically the R71G01 and dsx subsets (Fig. 5G)) drive slowing, which then enables the production of waggling followed by singing. While roles for P1 neurons, in solitary males, in both acute modulation of locomotion and persistent singing have been reported previously [5,12], through quantification of wing movements, we now additionally reveal that activation of particular subsets of P1 promotes a reliable behavioral sequence that includes a new behavior, waggling.

## Conclusion

*Drosophila* courtship has been characterized as a probabilistic sequence of distinct actions that unfold over minutes [2], with individual actions like courtship song being dynamically shaped by sensory cues on hundred-millisecond to second timescales [3,4]. Our findings reveal an intermediate level of organization: the stillness → waggling → singing sequence represents a stereotyped behavioral ‘chunk’, that is deployed within the otherwise variable flow of the courtship sequence. This suggests a hierarchical system where the nervous system can call upon structured sub-modules as needed, rather than generating every action de novo in response to external cues.

While we do not yet know if waggling serves a communicative function, our data strongly suggest a preparatory role. The dominant wing during waggling reliably predicts which wing the male will extend for subsequent singing, suggesting waggling acts as motor priming, a “warm-up” that facilitates the wing selection and extension required for courtship singing. The intermediate frequency of waggling (∼10.6 Hz), positioned between low-frequency thoracic vibrations and high-frequency song, further supports this preparatory function.

The mechanism underlying the P1-driven stillness → waggling → singing module contrasts with another recently described *Drosophila* courtship sequence. McKellar et al. [23] identified an “engagement motif” (proboscis extension, abdominal bending, and foreleg lifting) controlled by the aSP22 descending neuron pair, where increasing spike counts trigger cumulative, overlapping actions. Our sequence operates differently: actions are transitional rather than cumulative, and control originates from P1 in the central brain rather than from descending neurons. While we haven’t identified the specific neurons downstream of P1 that control waggling, activation of known song-production neurons (pIP10, pMP2, dPR1, TN1) [9,35–37] never elicited waggling (data not shown), indicating P1 coordinates distinct parallel pathways for waggling and singing.

Our results contribute to the growing body of evidence for functional heterogeneity within the P1 neuron population (Fig. 5G). The 11 P1 genotypes we tested do not represent distinct cell types but rather different, but overlapping subpopulations within the P1 neural space, allowing us to dissect functional domains. Comparing different driver line combinations revealed that R71G01-containing lines (splits 1, 2, 3, and r71_dsx) consistently induced the courtship sequence. For this subset, our results point to a clear split in the dsx-expressing and fru-expressing subsets: dsx+ P1 neurons drive the behavioral sequence, while fru+ P1 neurons do not. This molecular dissociation aligns with previous studies demonstrating functional specializations for dsx+ versus fru+ P1 neurons [32,33]. In addition, we find that Split6 (R15A01∩R22D03) induced the sequence without R71G01, suggesting a potentially parallel pathway operating independently of the R71G01-defined subset. In contrast, NP2631-intersection lines consistently failed to elicit the sequence.

Finally, P1 neurons appear to function as a context-dependent switch, orchestrating distinct motor programs based on the presence or absence of female motion cues. When females are moving, R71G01/P1 activation increases the gain of visual processing pathways to enable chasing and pursuit [38]. In contrast, our work shows that in the absence of such female motion cues, R71G01/P1 activation initiates an entirely different sequence: acute locomotor slowing, followed by waggling, and then a transition to song. This suggests P1 neurons translate the sensory context into adaptive actions, engaging a “chasing” program when the female is moving and a “slowing→waggling→singing” program when she is still. In both of these contexts, P1 neurons likely are ‘aware’ of the female’s presence through detection of pheromonal cues [32]. Once singing begins (following waggling), locomotor speed increases regardless of whether the sequence is naturally occurring (Fig. 2B) or optogenetically induced (Fig. 5C), readying the male for the next phase of the dynamic courtship interaction.

## STAR Methods

### Key resources table

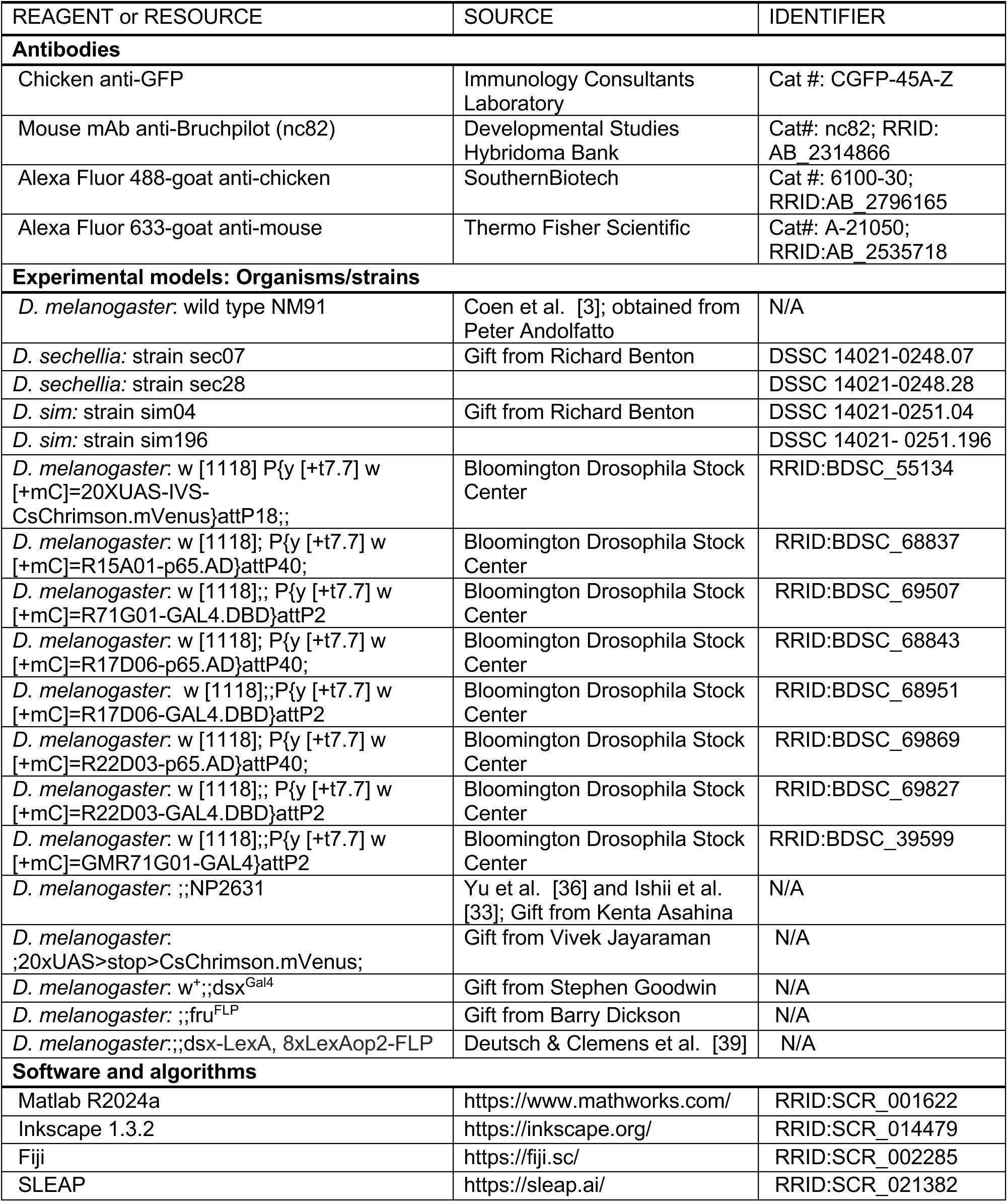

### Experimental model and subject details

Male *Drosophila melanogaster*, *D. simulans*, and *D. sechellia* flies were reared at 25°C on 12:12 light:dark cycles. *D. melanogaster* and *D. simulans* were maintained on standard cornmeal medium; *D. sechellia* on Formula 4-24 medium (Carolina) mixed with noni juice. Virgin males were collected within 8 hours of eclosion, aged 3–7 days, and single-housed except for optogenetic experiments (2–8 flies per vial). For courtship assays, individual virgin males were paired with age-matched virgin females that were also single-housed prior to experiments. For optogenetic experiments (expression patterns in Fig. S5), males were transferred to ATR-supplemented food (1mM) within 8 hours of eclosion. Genotype details are in the Key Resource Table and Table S1.

### Methods details

#### Courtship behavioral assays

Courtship interactions were recorded in a custom-fabricated behavioral chamber consisting of a 30 mm × 30 mm 3D-printed base (Formlabs Form 2, Black resin) and a clear vacuum-molded PETG dome. The arena was side-illuminated with 850 nm infrared LEDs, and videos were captured from above at 150 frames per second with 1,024 × 1,024 pixel resolution (30.3 pixels/mm) using a FLIR Blackfly S camera equipped with an 850 nm longpass filter. Acoustic signals were simultaneously recorded through microphones embedded in the chamber floor. Video files were compressed in real-time using GPU-accelerated H264 encoding to produce nearly lossless videos with independently seekable frames. This behavioral apparatus and recording protocol are described in detail in Pereira et al. (2022) [6]. Part of the wild-type *D. melanogaster* courtship dataset used in this study was previously published in Pereira et al. (2022) [6]. All *D. simulans* and *D. sechellia* data were newly collected for this manuscript.

Virgin males and females (3-7 days post-eclosion) were single-housed prior to experiments. All recordings began within 2 hours of incubator lights turning on to capture peak courtship activity. Individual male-female pairs were gently aspirated into the chamber and recorded for up to 30 minutes or until copulation occurred. For *D. sechellia* experiments, paper soaked with noni juice was placed under the chamber floor to provide species-appropriate olfactory cues. Only recordings containing courtship interactions were included in analyses; pairs showing no courtship behavior were excluded.

#### Optogenetics

To activate specific P1 neuron subsets, we used eleven genotypes expressing CsChrimson under different genetic drivers (see Table S1 for complete genotype list and Fig. S5 for anatomical expression patterns). Males were collected within 8 hours of eclosion and transferred to food supplemented with 1 mM all-trans-retinal (ATR) for at least three days prior to experiments. To prevent ATR degradation while maintaining circadian rhythms, vials were housed in custom-made blue acrylic light-filtering boxes.

Optogenetic experiments were performed using the same behavioral apparatus as courtship assays. Solitary males were individually placed in recording chambers and allowed to acclimate for 5 minutes before recording onset. Red light stimulation (627 nm) was delivered through programmable LEDs (Luxeon) mounted around the chamber.

The stimulation protocol consisted of 30-second trial blocks, each containing a 1-second light pulse followed by a 29-second inter-stimulus interval (Fig. 5B). Five LED intensities (1, 5, 25, 125, and 205 μW/mm²) were presented in randomized order, with each intensity repeated 10 times, resulting in a total experimental duration of 25 minutes per fly. Light intensity was calibrated using a power meter positioned at the chamber floor. Video and audio acquisition followed the same procedures as for courtship assays.

#### Anatomy

To visualize anatomical expression patterns of the eleven P1 neuron subsets, we performed whole-mount immunohistochemistry on adult male brains. Brains were dissected and stained following standard protocols (detailed protocols available at https://www.janelia.org/project-team/flylight/protocols). Primary antibodies included mouse anti-Bruchpilot (nc82, 1:30) for neuropil counterstaining and chicken anti-GFP (1:1000) to detect mVenus expression. Secondary antibodies were goat anti-chicken Alexa Fluor 488 (1:250) and goat anti-mouse Alexa Fluor 633 (1:250). All antibodies are listed in the Key Resources Table.

Serial optical sections were obtained at 1 μm intervals on a Zeiss 700 confocal microscope with a Plan-Apochromat 20×/0.8 NA objective. Confocal stacks were processed into maximum intensity projections using Fiji. For figure S5, original projections were paired with masked versions where non-specific expression outside the central brain was digitally removed to highlight P1 neuron populations. Figure 5G shows the masked anatomical patterns for clarity.

### Behavioral data analysis

#### Pose tracking

All videos were tracked using SLEAP [6], and then manually proofread. For each fly, we tracked four body parts: head, thorax, left wing tip, and right wing tip. The thorax position served as the primary reference point for fly location throughout the recording. Tracking data were exported at the original 150 fps temporal resolution for subsequent analyses. Tracked coordinates were first processed to reduce noise and tracking errors. Wing tip positions were smoothed by interpolating missing values using nearest-neighbor interpolation. Thorax coordinates underwent additional smoothing with a 3-frame moving average filter followed by a Savitzky-Golay filter.

#### Audio processing

Acoustic signals were recorded through 9 microphones embedded in the chamber floor and digitized at 10 kHz for courtship assays and 5 kHz for optogenetic experiments. Audio channels were first baseline-subtracted then passed through a band-pass filter (50-1000 Hz). The three channels with highest amplitude at each time point were selected and averaged to produce a single audio trace. Audio and video streams were synchronized post-hoc using camera shutter signals recorded.

#### Feature extraction

The body axis for each fly was defined as the vector from thorax to head. Wing angles were calculated as the angle between each wing (thorax to wing tip vector) and the body axis, with 0° representing wings folded against the body and 90° representing full extension. The dominant wing was defined as the wing with greater mean amplitude throughout each bout.

Locomotor metrics were derived from thorax positions. Speed was calculated as the displacement along the heading direction between consecutive frames. All speed traces shown in figures were smoothed with a median filter. For z-scored speeds shown in figures, z-scoring was performed on the entire recording for each animal.

To quantify male-female interactions, we computed three key spatial relationships (Fig. 1H): (1) the Euclidean distance between male and female thoraxes, (2) the body angle between the flies’ body axes, and (3) the target angle between the male’s body axis and the vector pointing from male to female thorax. For spatial visualization, we generated female-centric plots. The location heatmap (Fig. 1I) was created by converting male positions relative to the female into Cartesian coordinates based on thorax-to-thorax distance and the angle from the female’s body axis. These positions were binned into a hexagonal grid (0.10 mm edge length). The orientation quiver plot (Fig. 1J) displayed male body orientations at different positions around the female, with data grouped into 14 × 14 spatial bins (approximately 0.86 mm per bin). Each bin’s vector base represents the average male position, its direction shows the average relative body angle, and its length scales with the number of observations in that bin.

#### Waggle and song bout detection

Waggling and courtship song bouts were identified based on distinct kinematic and acoustic signatures. For courtship song bouts detection, we used a custom written script that identified periods of substantial wing extension accompanied by elevated acoustic amplitude. Bout boundaries were defined as the frames where wing angles exceeded and subsequently returned to baseline. All detected song bouts were manually proofread. For waggling detection, different approaches were used depending on the dataset. In courtship assay datasets (*D. melanogaster*, *D. simulans*, and *D. sechellia*), waggle bouts were manually annotated by identifying periods of rhythmic, anti-phase wing movements lacking acoustic signals. For optogenetic experiments, we used a custom written waggling detector. Wing angles were band-pass filtered (8-18 Hz), and the Hilbert transform was applied to extract instantaneous phase and amplitude. Waggling was identified when wings oscillated in anti-phase with sufficient amplitude above baseline. Waggle bout boundary was set from the onset to the offset of the continuous oscillatory wing movement. Wing oscillations must persist for at least 3 cycles to be counted as waggling. The automated detector achieved 89.5% precision and 83.6% recall when validated against manually annotated data. All automatically detected waggling bouts were subsequently manually proofread.

#### Stillness definition

Stillness was defined as periods when both male and female flies remained nearly stationary. To determine appropriate speed thresholds, we applied a knee-point algorithm to the cumulative distribution of speeds pooled across all recordings. This algorithm identifies the point of maximum curvature by calculating the perpendicular distance from each point on the normalized CDF to a line connecting the distribution endpoints. The resulting inflection points distinguished stationary from mobile behavior: 0.67 mm/s for males and 0.30 mm/s for females (Fig. S2). Periods were classified as stillness only when both flies simultaneously maintained speeds below their respective thresholds.

#### Temporal linking of waggle bouts

To examine coupling between waggling and singing, we measured the gap duration from each waggle bout offset to the subsequent song bout onset (Fig. 3A). We classified transitions as “linked” or “unlinked” using k-means clustering (k=2) on log-transformed gap durations, with 0.15 seconds as the dividing threshold. Waggle bouts not followed by song within the recording period were excluded from this analysis.

#### Behavioral state transitions

Four mutually exclusive behavioral states were defined: waggling, singing, stillness, and other. Each frame of the recording was assigned to one of these states based on a hierarchical classification scheme. First, stillness frames were identified based on the speed criteria described above. Next, waggling and singing bouts were assigned to their respective frames, overwriting any stillness assignments. This ensured that periods where flies were stationary while waggling or singing were correctly classified by their active behavior rather than as stillness. All remaining frames were classified as “other.”

To focus analysis on meaningful behavioral periods, minimum bout duration thresholds were established for each state. For stillness and other states, bout duration thresholds were determined using the knee-point algorithm applied to the distribution of bout durations, yielding thresholds of 0.71 s and 0.72 s, respectively (Fig. S4). Waggling and singing bouts retained their detection-based durations as described above. Bouts shorter than these thresholds were excluded from subsequent analyses.

State transition probabilities were computed by tallying all bout-to-bout transitions across recordings. To assess statistical significance, we performed permutation tests (10,000 iterations) where bout labels were randomly shuffled within each recording while preserving the frequency and duration of each state. This null model tested whether observed transition probabilities exceeded chance expectations (one-tailed test for enrichment). P-values were corrected for multiple comparisons using the Benjamini-Hochberg false discovery rate (FDR) procedure. Figure 4 presents the 3×3 transition probability submatrix for the three courtship-relevant states (stillness, waggling, singing), while the complete 4×4 transition matrix including all transitions to and from “other” states is shown in Fig. S4. For three-gram analysis, we examined all sequences of three consecutive behavioral bouts, excluding sequences containing “other” states or self-transitions.

### Quantification and statistical analysis

All statistical analyses were performed using custom scripts in MATLAB R2024a. Statistical tests were selected based on data distribution and experimental design. The Shapiro-Wilk test was used to assess normality of data distributions, which determined whether parametric or non-parametric tests were applied.

Tests used in this study included: paired t-tests for normally distributed paired comparisons; Wilcoxon signed-rank tests for non-normally distributed paired data; one-way ANOVA with Tukey’s post-hoc correction for normally distributed data across multiple groups; Kruskal-Wallis tests with Dunn-Šidák post-hoc correction for non-normally distributed multi-group comparisons; Friedman tests for non-parametric repeated measures analysis when comparing multiple conditions within subjects; and one-tailed permutation tests (10,000 iterations) with Benjamini-Hochberg false discovery rate correction for state transition probabilities.

Statistical details for each comparison are provided in figure legends, including sample size, test used, and p-values. Data are presented as mean ± SEM for bout-triggered averages and time series. Box plots display median (center line), interquartile range (box), and 1.5×IQR (whiskers), with individual data points overlaid. Violin plots show full data distributions with median and quartiles marked. Significance levels are indicated as ns (not significant), * (p < 0.05), ** (p < 0.01), or *** (p < 0.001).

## Supplemental information

Document S1. Figures S1–S8, Tables S1, and supplemental references

Video S1. A male *Drosophila melanogaster* performs waggling towards a female, related to figure 1. The video was recorded at 150 frames per second (fps) and is played back at 25 fps.

Video S2. A male *Drosophila melanogaster* performs courtship singing towards a female, related to figure 1. The video was recorded at 150 frames per second (fps) and is played back at 25 fps.

Video S3. A male *Drosophila simulans* performs waggling towards a female, related to figure S1. The video was recorded at 150 frames per second (fps) and is played back at 25 fps.

Video S4. A male *Drosophila sechellia* performs waggling towards a female, related to figure S1. The video was recorded at 150 frames per second (fps) and is played back at 25 fps.

Video S5. The stereotyped stillness-waggling-singing courtship sequence in *Drosophila melanogaster*, related to figure 4. The sequence begins as both flies decelerate to a state of mutual stillness. The male then initiates waggling followed by song. The video was recorded at 150 frames per second (fps) and is played back at 25 fps.

## Resource availability

### Lead contact

Further information and requests for resources and reagents should be directed to the lead contact, Mala Murthy.

### Materials availability

This study did not generate new transgenic reagents. Fly strains and lines used in this study are listed in the key resources table.

### Data and code availability

Data reported in this paper will be shared by the lead contact upon request. Code for detecting and annotating waggle bouts is on Github at github.com/murthylab and will be publicly available upon publication. Any additional information required to reanalyze the data reported in this paper is available from the Lead Contact upon request.

## Acknowledgments

We thank Yoon Woo Park for help with data annotation. We thank Junyu Li and Talmo Pereira [6] for collecting and proofreading the wild type *D. melanogaster* courtship data. We thank Richard Benton, Kenta Asahina, Vivek Jayaraman, Barry Dickson, and Stephen Goodwin for gifting flies. We thank Jan Clemens and the entire Murthy lab for helpful discussions. This research was supported by the following grants to MM: HHMI Faculty Scholar award, NIH NINDS R35 Research Program Award, and NIH BRAIN R01 NS104899.

## Author contributions

Conceptualization, X.L., K.T., and M.M.; methodology, X.L., K.T., and Y.G.; data collection, X.L., K.T., and Y.G.; investigation, X.L., K.T., Y.G., and M.M.; data curation, X.L., K.T.; formal analysis, X.L.; visualization, X.L.; supervision and acquisition of funding: M.M.; writing—original draft, X.L. and M.M.; writing—review & editing, X.L., K.T., and M.M.

## Declaration of interests

The authors declare no competing interests.

---

**Figure S1 - Related to Figure 1.**
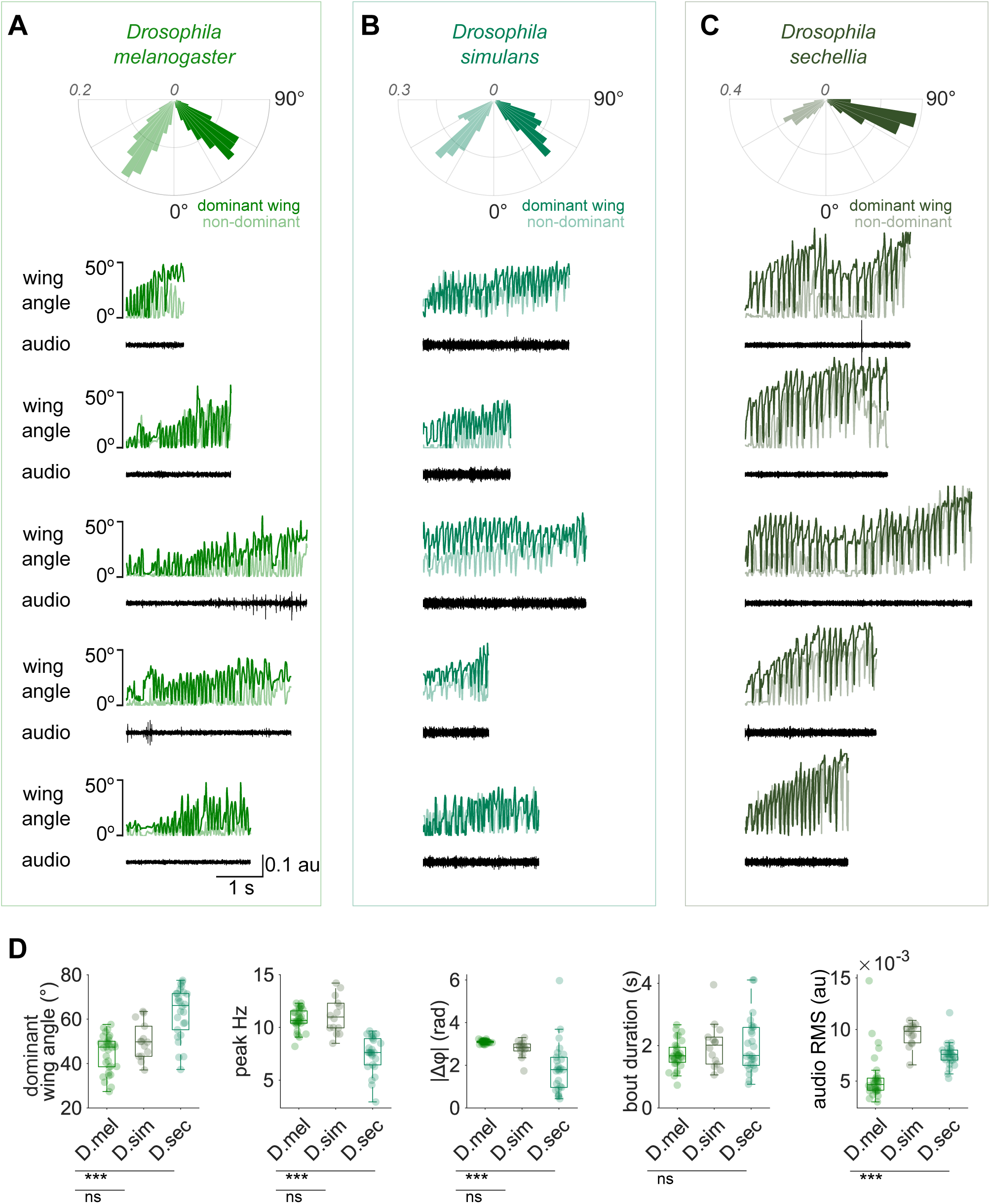
*Drosophila simulans and Drosophila sechellia* males also produce wing waggling during courtship. A-C. Top: Polar histograms show the distribution of maximum wing extension angles for the dominant (darker shade) and non-dominant (lighter shade) wings during waggling in each species. The dominant wing is defined as the side with the greater mean amplitude over the waggling bout. Bottom: Example traces from single waggling bouts in each species, showing wing angles and simultaneously recorded microphone audio. All traces share the same scale bar. D. Species comparison of waggling metrics. Box plots show median ± IQR, whiskers extend to 1.5×IQR, dots mark individual per-fly medians. *D. sechellia* extends its wings wider than either *D. melanogaster* or *D. simulans* (Kruskal–Wallis with Dunn–Šidák post-hoc, ***p < 0.001) and has a lower peak waggling frequency (ANOVA with Tukey post-hoc, ***p < 0.001). *D. melanogaster* and *D. simulans* exhibit nearly anti-phase wing movements during waggling, whereas *D. sechellia* shows weaker phase locking (ANOVA with Tukey post-hoc, ***p < 0.001). Bout durations are comparable across species (ANOVA, p = 0.20). Audio amplitude during waggling differs significantly between species (ANOVA, ***p < 0.001) but remains much quieter than courtship song (see Figure 1G). N = 32 flies/1,857 bouts (*D. melanogaster*), 14 flies/174 bouts (*D. simulans*), and 26 flies/528 bouts (*D. sechellia*).

**Figure S2 - Related to Figure 2.**
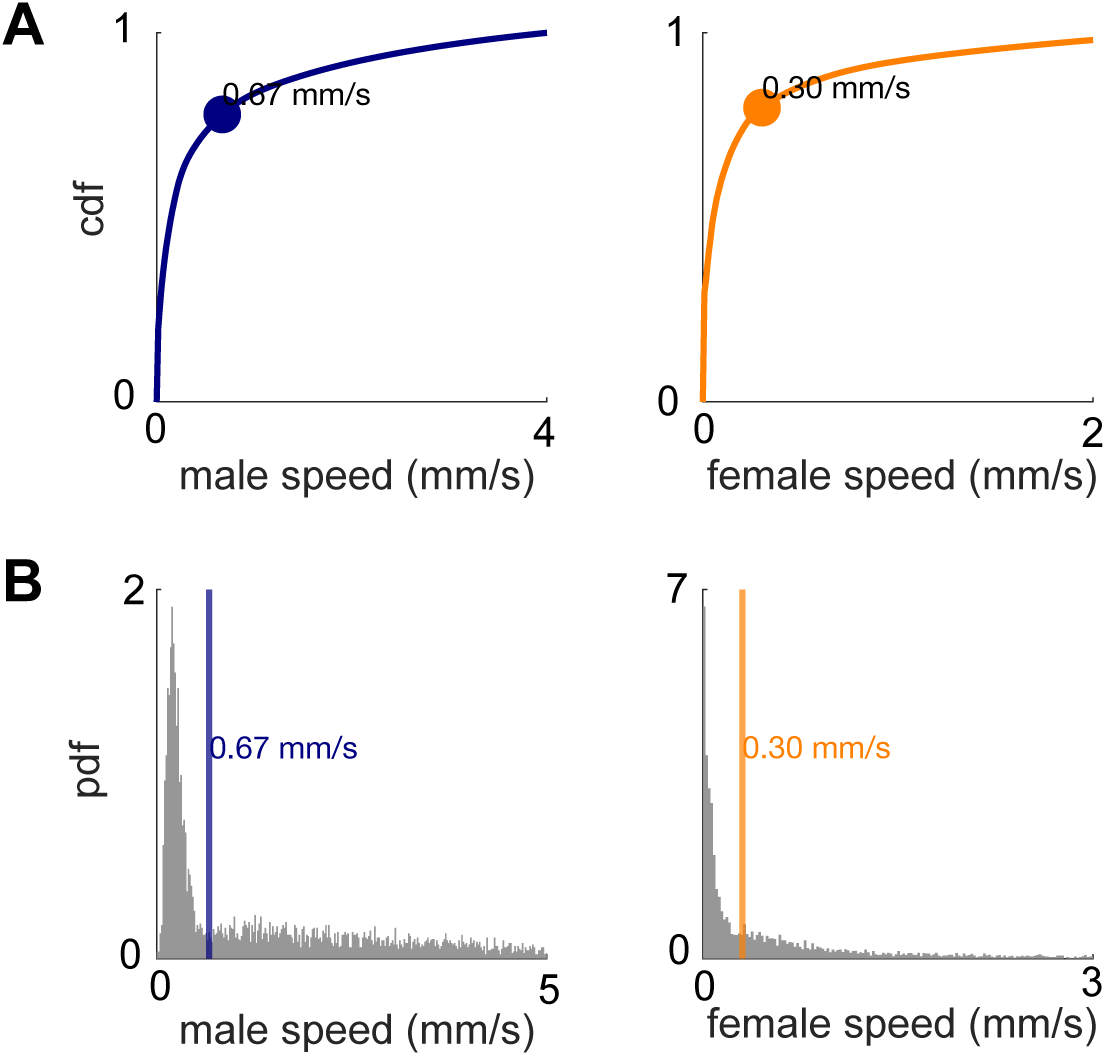
Defining the “stillness” threshold based on male and female locomotion. A. Cumulative probability distributions (CDFs) of male (navy) and female (orange) speeds across all frames. Dots mark the knee points (inflection points) at 0.67 mm/s for males and 0.30 mm/s for females. These thresholds define the “stillness” threshold, periods when both the male and female remain nearly stationary. B. Probability density functions (PDFs) of male (left) and female (right) speeds (gray) within waggle and song bouts combined. Vertical lines mark the corresponding stillness thresholds.

**Figure S3 - Related to Figure 3.**
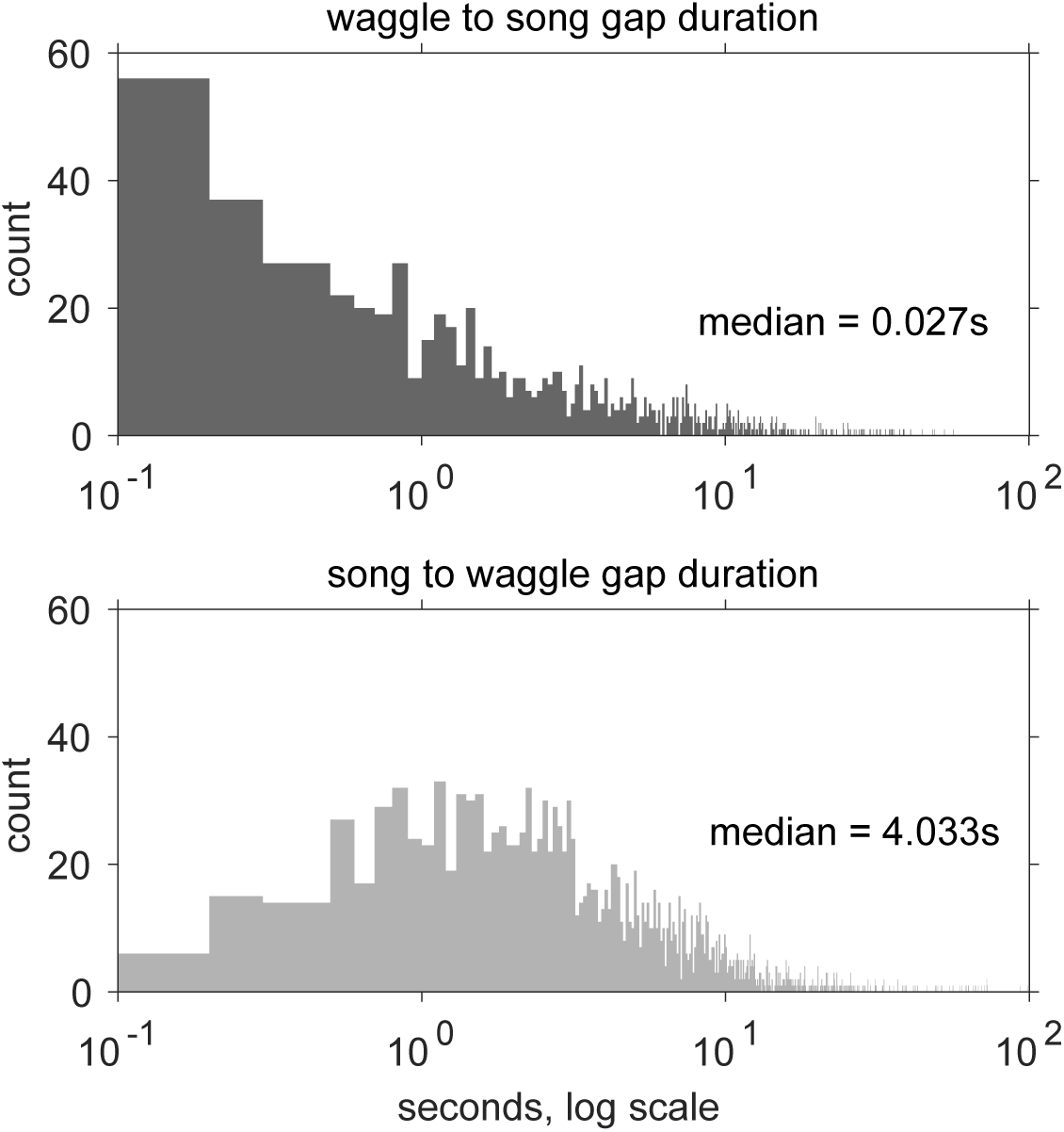
Waggling is followed by, not preceded by, singing. Top panel: Distribution of gap durations between the end of waggle bouts and the start of the subsequent song bout. Bottom panel: Distribution of gap durations between the start of waggle bouts and the end of the previous song bout.

**Figure S4 - Related to Figure 4.**
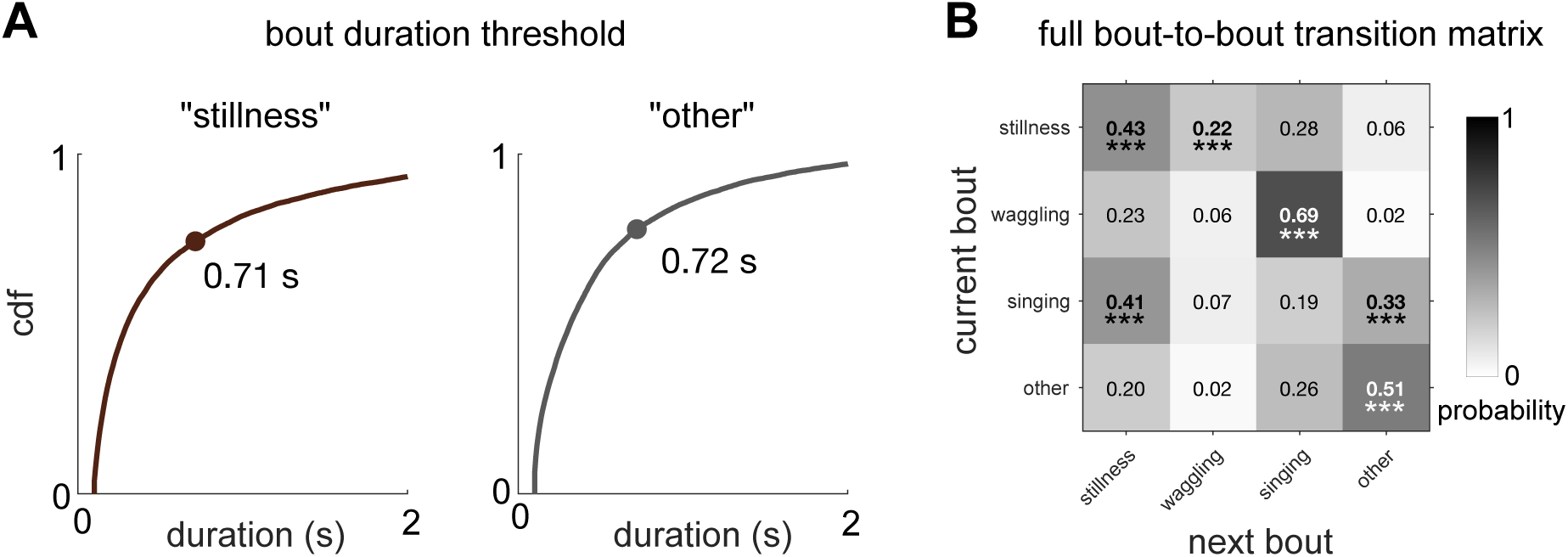
Bout duration thresholds and full behavioral transition matrix. A. Cumulative probability distributions of durations for “stillness” (left) and “other” (right) states. Dots mark the duration thresholds used to define bouts, identified using the knee-point algorithm (see Methods). B. Complete four-state bout-to-bout transition probability matrix. Transition probabilities between all behavioral states (stillness, waggling, singing, and other). Statistically significant transitions are highlighted (one-tailed, permutation test, ***p < 0.001). This complements the three-state submatrix shown in Figure 4B.

**Figure S5 - Related to Figure 5.**
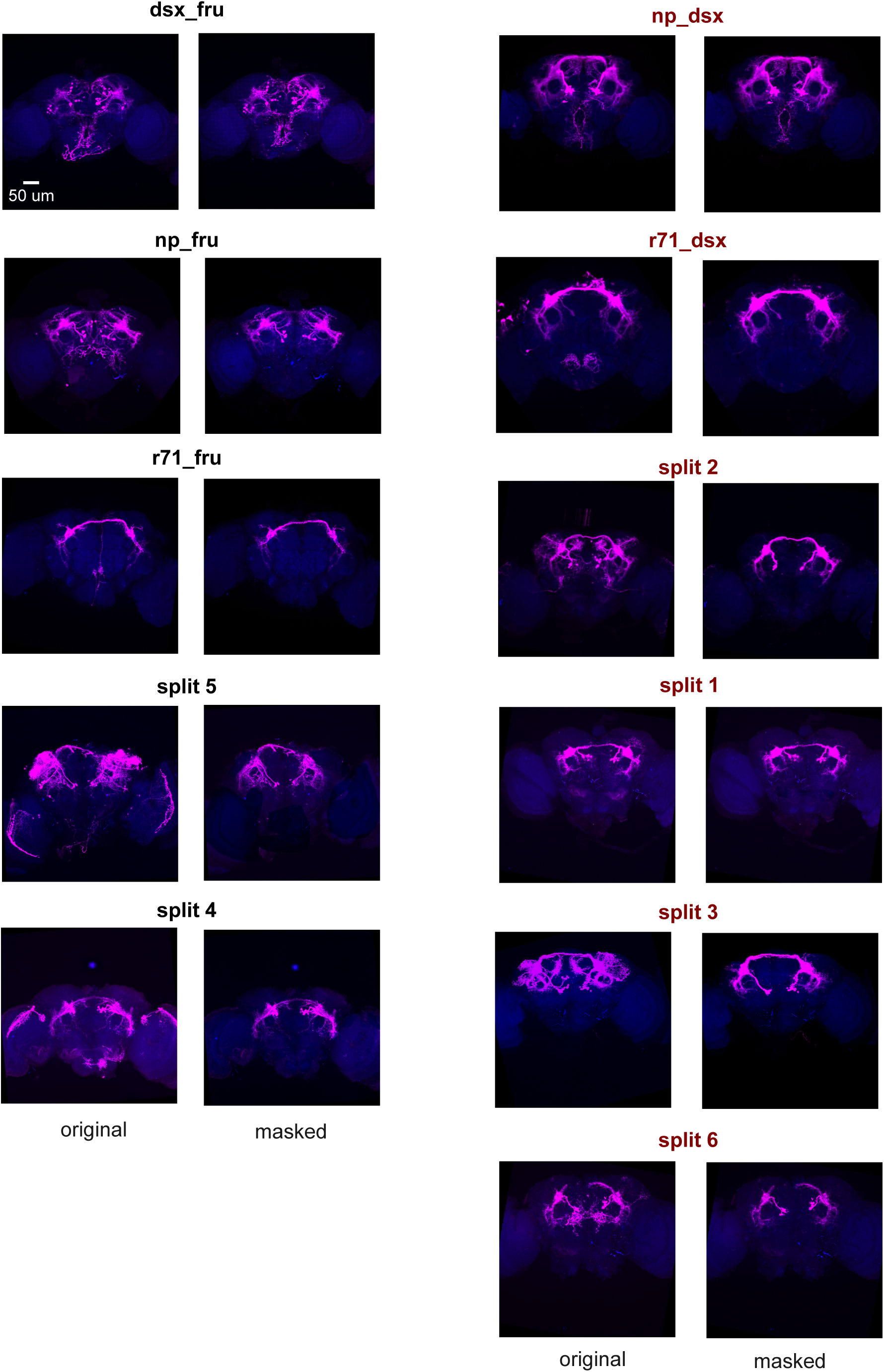
Anatomical expression of P1 neuron subsets. Maximum intensityprojections of confocal stacks showing brain expression patterns for all 11 genotypes used in optogenetic experiments (magenta: anti-GFP; blue: neuropil counterstain). Each pair shows original (left) and masked (right) images; masked images have non-P1 expression removed for clarity. While most driver combinations target mostly P1 neurons, some additional neurons are labeled in each line.

**Figure S6 - Related to Figure 5.**
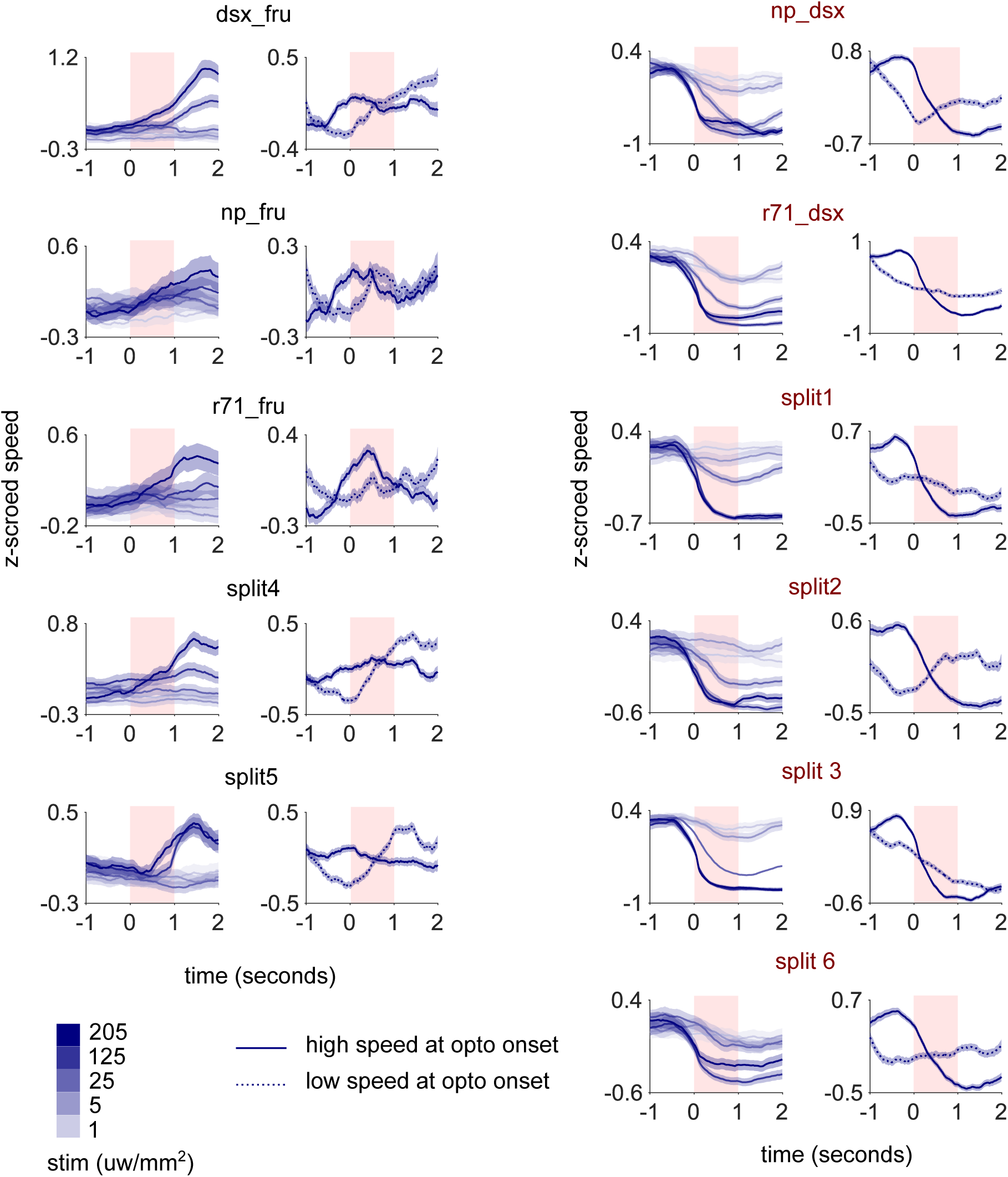
Speed dynamics following optogenetic activation of P1 neuron subsets. Z-scored walking speed aligned to optogenetic onset (red shaded area), sorted by genotype. For each genotype, left panels show responses split by optogenetic stimulus strength (5 levels, light to dark blue); right panels show responses split by initial locomotor state at stimulation onset (high speed: solid lines; low speed: dashed lines). High/low speed groups are determined by median split of speed at optogenetic onset across all stimulus levels. N = 17 (dsx_fru), 15 (np_dsx), 14 (np_fru), 19 (r71_dsx), 15 (r71_fru), 19 (split1/split P1a), 18 (split2), 19 (split3), 19 (split4), 19 (split5), 18 (split6) males.

**Figure S7 - Related to Figure 5.**
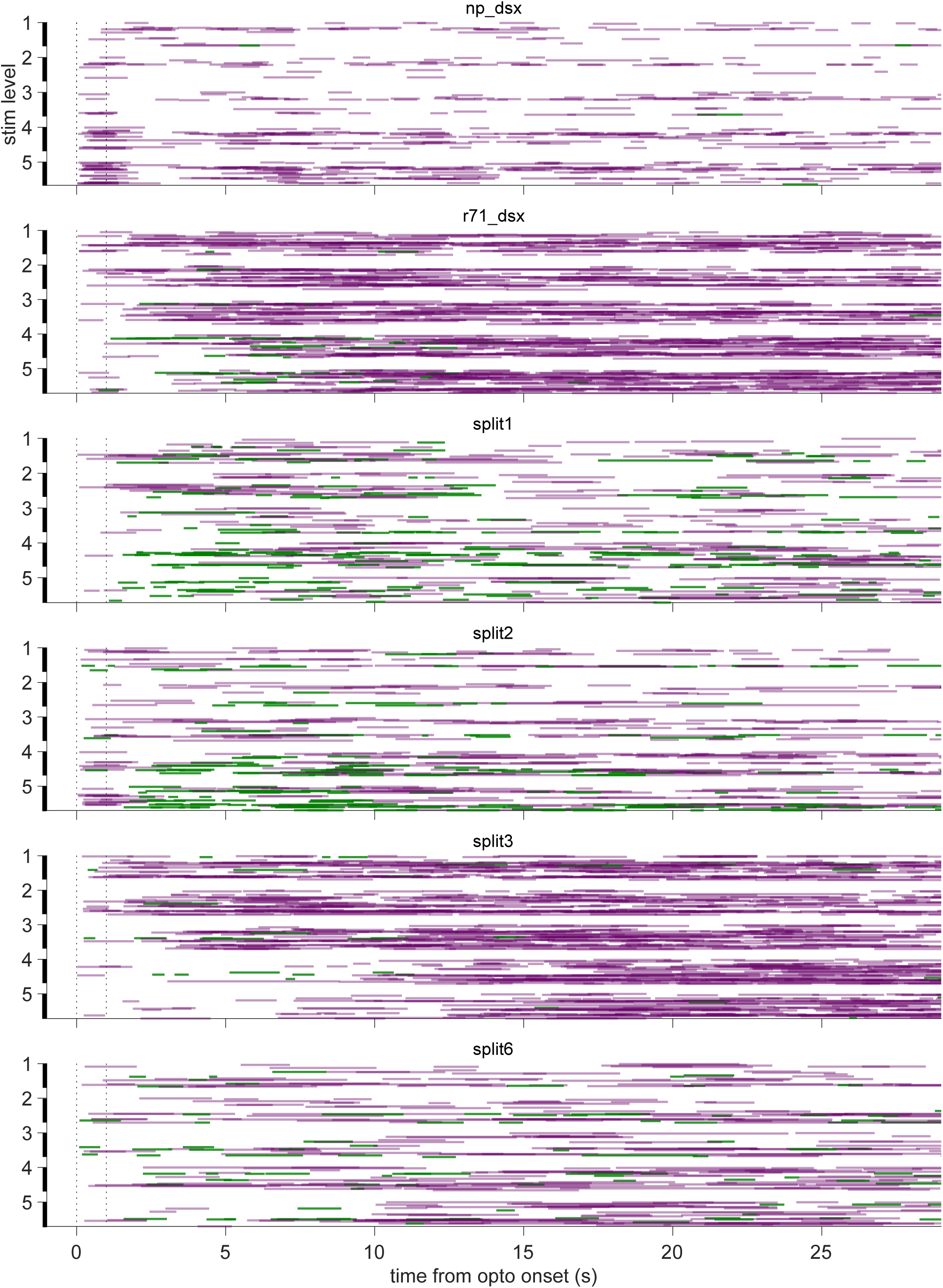
Raster plots of optogenetically evoked waggling and singing. Behavioral responses for each effective genotype following optogenetic stimulation. Each row represents a single trial, with horizontal bars indicating detected waggle (green) and song (purple) bouts. Dashed lines represent onset and offset of optogenetic stimulation. Trials are grouped by stimulation strength (levels 1-5: 1, 5, 25, 125, 205 μW/mm²). N = 15 (np_dsx), 19 (r71_dsx), 19 (split1/split P1a), 18 (split2), 19 (split3), 18 (split6) males.

**Figure S8 - Related to Figure 5.**
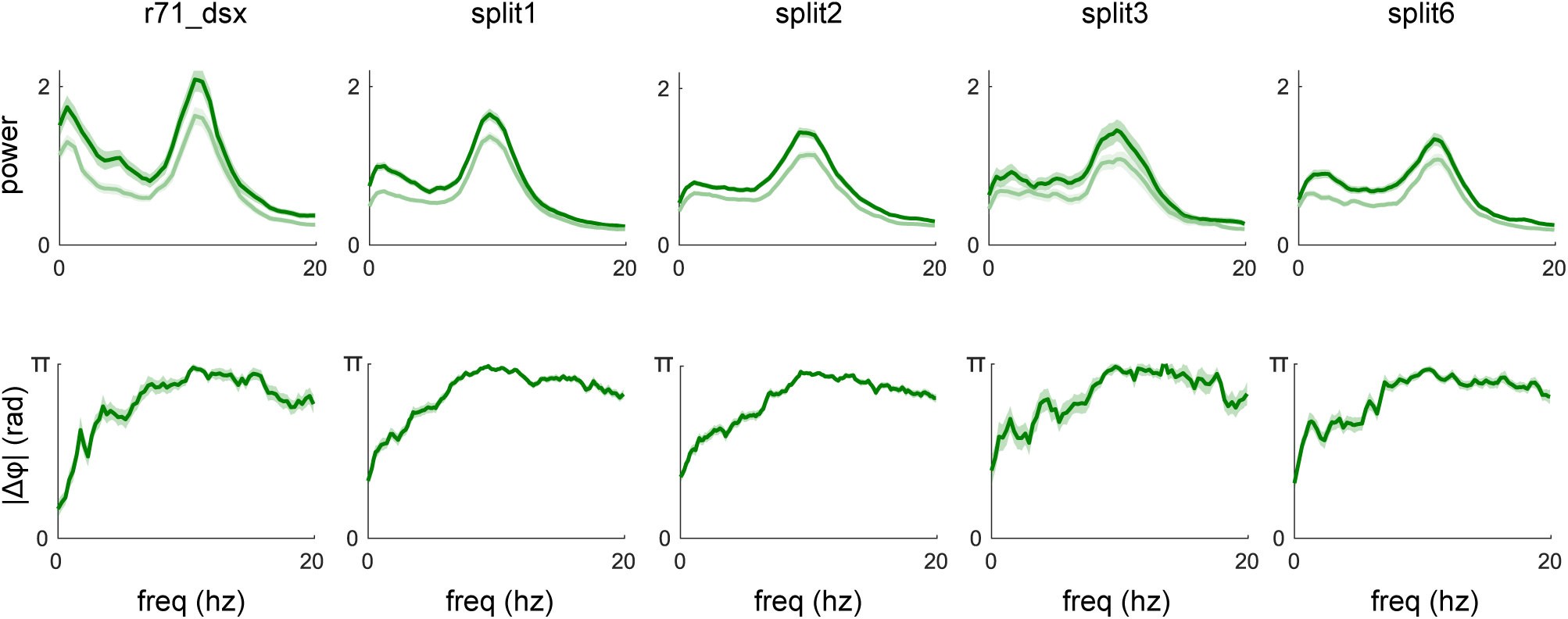
Wing movement characteristics of optogenetically evoked waggling. Power spectra (top) and absolute inter-wing phase difference |Δφ| (bottom) during optogenetically evoked waggling bouts for each effective genotype (mean ± SEM). N = 19 (r71_dsx), 19 (split1/split P1a), 18 (split2), 19 (split3), 18 (split6) males.

**Table S1- Related to Figure 5.** Genotypes of P1 driver lines used for optogenetic experiments.

| abbreviation<br>used in this<br>manuscript | genotype | reference |
| --- | --- | --- |
| split1 | UAS-CsChrimson.mVenus;<br>R15A01.AD/+; R71G01.DBD/+ | Hoopfer et al.[1], as P1<sup>a</sup> |
| split2 | UAS-CsChrimson.mVenus;<br>R17D06.AD/+; R71G01.DBD/+ | Zhang et al.[2], as P1<sup>c</sup> |
| split3 | UAS-CsChrimson.mVenus;<br>R22D03.AD/+; R71G01.DBD/+ |  |
| split4 | UAS-CsChrimson.mVenus;<br>R15A01.AD/+; R17D06.DBD/+ |  |
| split5 | UAS-CsChrimson.mVenus;<br>R22D03.AD/+; R17D06.DBD/+ | Zhang et al.[2], same combination as P1<sup>d</sup> |
| split6 | UAS-CsChrimson.mVenus;<br>R15A01.AD/+; R22D03.DBD/+ | Zhang et al.[2], as P1<sup>b</sup> |
| dsx_fru | ;UAS>stop> CsChrimson.mVenus/+;<br>dsx<sup>Gal4</sup>/ Fru<sup>FLP</sup> | Ishii et al.[3], similar to dsx<sup>GAL4</sup>, UAS > stop<br>> CsChrimson, fru<sup>FLP</sup> |
| r71_fru | ;UAS>stop> CsChrimson.mVenus/+ ;<br>Fru<sup>FLP</sup>/R71G01 | Coleman et al.[4], similar to w; 71G01-<br>p65.AD/+;UAS-CsChrimson, fru-DBD/+ + |
| r71_dsx | +/UAS>stop> CsChrimson.mVenus; dsx-LexA, 8xLexAop2-FLP/R71G01 | Coleman et al.[4], similar to w; 71G01-<br>p65.AD/UAS-CsChrimson; dsx-DBD/+ |
| np_fru | ;NP2631/UAS>stop><br>CsChrimson.mVenus; Fru<sup>FLP</sup>/+ |  |
| np_dsx | NP2631/UAS>stop><br>CsChrimson.mVenus; dsx-LexA,<br>8xLexAop2-FLP/+ | Ishii et al.[3], similar to NP2631, UAS ><br>stop > CsChrimson, dsx<sup>FLP</sup> |

## Notes

### Competing Interest Statement

The authors have declared no competing interest.

